# A mediterranean-mimicking diet harnesses gut microbiota–derived 3-IAA to rejuvenate T cell

**DOI:** 10.64898/2026.01.05.697620

**Authors:** Xin Yu, Wenge Li, Hongfang Feng, Zhiyu Li, Hongmei Zheng, Shengrong Sun, Juanjuan Li, Bei Li, Qi Wu

## Abstract

Diet profoundly shapes the tumor microenvironment through metabolite modulation, yet the mechanisms linking diet, gut microbiota, and antitumor immunity remain unclear. Here, we demonstrate that a mediterranean-mimicking diet (MedDiet) suppresses tumor growth by orchestrating a microbiota–metabolite–immune axis. MedDiet selectively enriched *Bacteroides thetaiotaomicron* (*B. thetaiotaomicron*) in the gut, whose administration recapitulated tumor suppression. Both MedDiet and *B. thetaiotaomicron* elevated plasma levels of the tryptophan metabolite indole-3-acetic acid (3-IAA). Mechanistically, 3-IAA enhanced CD8^+^ T cell cytotoxicity and alleviated exhaustion by inhibiting the integrated stress response, sustaining antitumor immunity *in vivo*. Importantly, 3-IAA synergized with anti–PD-1 therapy to further restrain tumor progression. Finally, we developed a MedDiet Score to predict patient survival and response to immunotherapy. These findings reveal that dietary modulation of gut microbiota can reshape systemic metabolites to potentiate T cell–mediated antitumor responses. Our study identifies 3-IAA as a metabolite-based adjuvant for immune checkpoint therapy and highlights the translational potential of diet-driven immunomodulation in cancer treatment.

## Introduction

Malignant neoplasms emerge from the complex interplay of genetic predisposition, environmental exposures, and modifiable lifestyle factors, among which diet represents a particularly powerful determinant. Epidemiological analyses estimate that dietary patterns contribute to nearly one-third of cancer cases(Irigaray et al., 2007; Wu et al., 2022). However, the mechanistic pathways linking nutrient intake to tumor initiation and progression remain incompletely understood. Disentangling these connections has been challenging due to heterogeneity in dietary assessments, inter-individual variation in nutrient metabolism, and the extensive biotransformation of dietary components by the gut microbiota.

Emerging studies converge on the idea that diet influences tumor biology not only through systemic metabolic alterations but also by remodeling the gut microbiome, which generates bioactive metabolites that govern tumor–immune interactions. For example, ketogenic feeding elevates the ketone body β-hydroxybutyrate, which constrains colorectal tumorigenesis by attenuating epithelial proliferation via the hydroxycarboxylic acid receptor 2 (HCAR2)–homeodomain-only protein homeobox (HOPX) axis(Dmitrieva-Posocco et al., 2022). Caloric restriction enriches *Bifidobacterium* species, whose acetate production enhances CD8⁺ T cell interferon-γ secretion and antitumor cytotoxicity(Mao et al., 2023). Conversely, high-fat diets dominated by butter promote long-chain acylcarnitine accumulation, impairing mitochondrial integrity and effector function in NK and CD8⁺ T cells(Kunkemoeller et al., 2025). These studies underscore that not only the quantity but also the qualitative source of dietary nutrients critically determines immune competence within the tumor microenvironment. Against this backdrop, the Mediterranean diet (MedDiet)—a plant-forward dietary pattern characterized by abundant intake of vegetables, fruits, whole grains, legumes, nuts, and olive oil, along with moderate fish, poultry, and dairy consumption——has drawn substantial attention for its broad health benefits(Bolte et al., 2023; Li et al., 2020; Liu et al., 2025a). Integrating dietary, metabolomic, and clinical data from the Spanish PREDIMED trial(Estruch et al., 2018) and three U.S. prospective cohorts (the Nurses’ Health Study (NHS), NHSII, and Health Professionals Follow-Up Study [HPFS])(Li et al., 2020; Liu et al., 2025b), investigators identified and validated a comprehensive metabolic signature associated with adherence to the MedDiet. This consistent metabolic signature encompassed 45 lipid species and acylcarnitines (20% of all assayed lipids), 19 amino acids (representing 44% of all analyzed amino acids), 2 vitamins (29% of assayed vitamins/cofactors), and 1 xenobiotic compound. The selected metabolites demonstrated strong, consistent associations with MedDiet adherence across datasets. Notably, among the 67 metabolites constituting the core signature, those linked to fish and seafood intake—particularly highly unsaturated lipid species—showed the greatest reproducibility, underscoring the pivotal contribution of marine-derived lipids to MedDiet adherence(Li et al., 2020; Liu et al., 2025a). Extensive clinical and epidemiological evidence links adherence to the MedDiet with reduced incidence of cardiovascular disease(Li et al., 2020), type 2 diabetes(Salas-Salvadó et al., 2011), cancer(Bolte et al., 2023), and neurodegeneration(Liu et al., 2025a), alongside improved inflammatory and metabolic profiles. These convergent findings establish the MedDiet as one of the most sustainable and health-promoting dietary patterns worldwide. However, despite its well-documented clinical benefits, the molecular dialogue between the MedDiet, microbiota-derived metabolites, and T cell stress-adaptive pathways remains poorly understood. Elucidating this mechanistic interface may provide key insights into how nutritional inputs recalibrate immune homeostasis and influence tumor immunosurveillance.

The integrated stress response (ISR) represents a conserved signaling hub that restrains global protein synthesis under conditions of metabolic stress, hypoxia, infection, and oxidative stress (Costa-Mattioli and Walter, 2020; Pakos-Zebrucka et al., 2016; Yu et al., 2025). Central to this pathway is phosphorylation of eukaryotic initiation factor 2α (eIF2α), which suppresses global translation while activating transcription factors such as activating transcription factor 4 (ATF4) and C/EBP homologous protein (CHOP) to govern cell fate decisions (Costa-Mattioli and Walter, 2020). Mounting evidence indicates that ISR activation not only accelerates malignant progression by enhancing tumor cell proliferation, metastasis, and therapy resistance but also imposes immunosuppressive constraints within the tumor microenvironment(Cao et al., 2024, 2019). Within tumors, ISR activation in CD8⁺ T cells drives functional exhaustion, diminishes interferon-γ secretion, and compromises immunosurveillance(Cao et al., 2019). How diet and microbial metabolites intersect with the ISR to shape CD8⁺ T cell fate remains a critical unresolved question.

Here, we identify a diet–microbiome–metabolite axis that restores CD8⁺ T cell function through suppression of the ISR. We show that MedDiet, enriched in fiber and fish oil, selectively expands *Bacteroides thetaiotaomicron* (*B. thetaiotaomicron*), which catalyzes the conversion of tryptophan into the indole derivative indole-3-acetic acid (3-IAA). Elevated systemic 3-IAA alleviates ISR-driven translational repression in CD8⁺ T cells, thereby mitigating exhaustion and preserving cytotoxicity. Functionally, administration of 3-IAA phenocopied the effects of MedDiet and microbial enrichment and synergized with anti–PD-1 therapy to further restrain tumor progression. These findings position 3-IAA as a metabolite-based immunotherapeutic adjuvant that calibrate T cell competence and highlights microbiome-targeted nutritional strategies as a promising frontier for enhancing immunotherapy and advancing cancer prevention.

## Result

### MedDiet suppresses tumor growth across various murine models

To experimentally recapitulate the essential features of MedDiet, we formulated a series of isocaloric mouse diets with precisely defined macronutrient sources. Each diet maintained a constant protein content and caloric density while differing in lipid composition and fiber enrichment. Specifically, the lipid component was systematically varied to include olive oil, soybean oil, fish oil, or palm oil as the primary fat source, allowing us to isolate the effects of dietary lipid quality and fiber content on tumor progression while minimizing potential confounders such as protein intake or caloric load (Fig. 1A). In male mice bearing subcutaneous MC38 colon carcinoma, only the MedDiet—distinguished by its high fiber and fish oil content—produced a robust suppression of tumor growth compared with all other diets (Fig. 1B–C). This effect occurred without measurable changes in body weight, food consumption, or fasting glucose, excluding systemic metabolic toxicity as a confounding factor. To assess the reproducibility and generalizability of this effect, we extended our analyses to additional tumor models. Strikingly, MedDiet-fed mice also exhibited pronounced growth inhibition of MCA205 fibrosarcoma (Fig. 1D), demonstrating that the tumor-suppressive effect was not model-specific, but rather represented a consistent biological response. These findings suggest that MedDiet restrains tumor progression not through global metabolic alterations but likely through the modulation of tumor-intrinsic signaling pathways or the host immune microenvironment.

**Figure 1.**
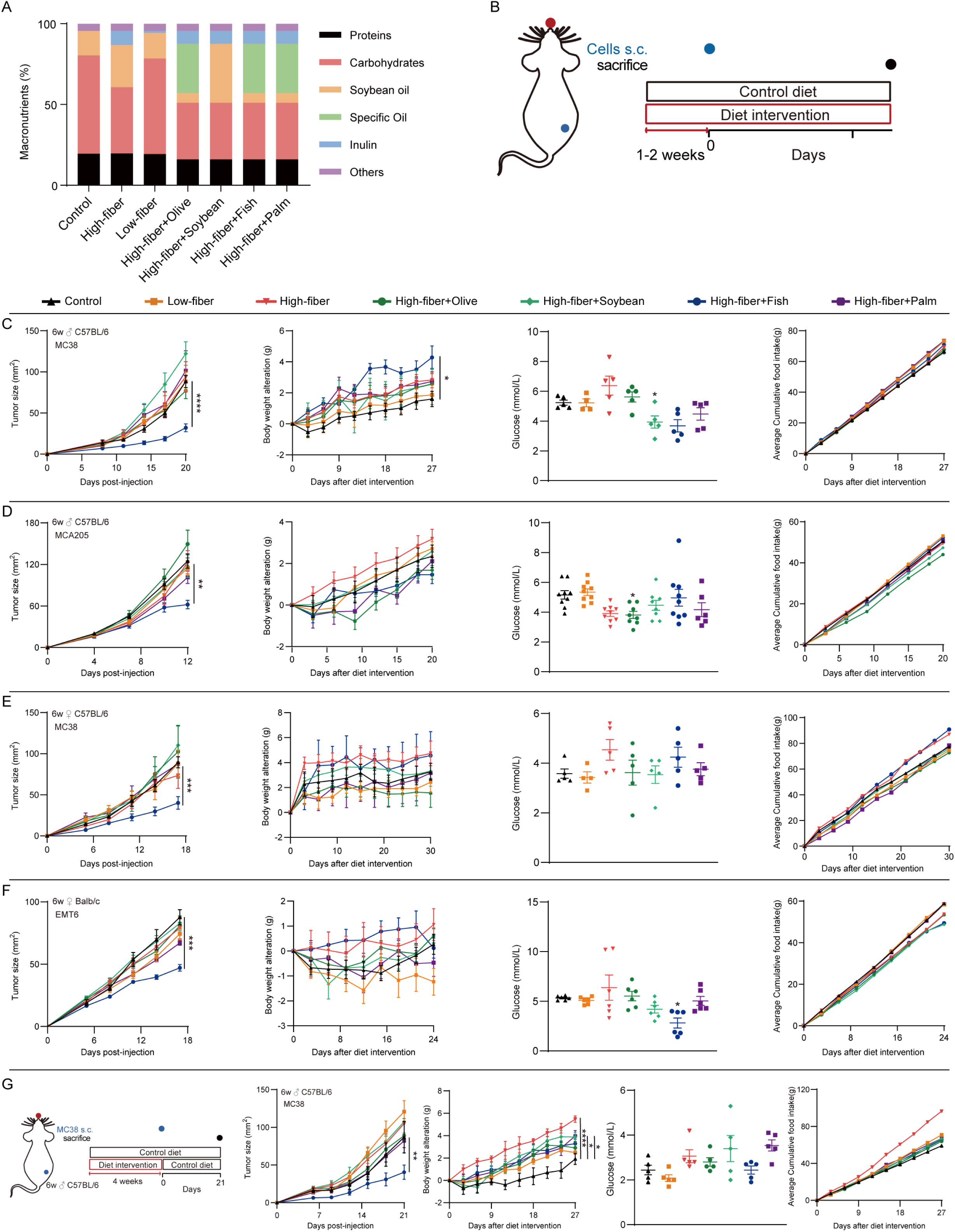
MedDiet inhibits tumor progression. (A) Composition of different dietary regimens. (B–F) Subcutaneous tumor models subjected to dietary interventions: (B) Schematic of the experimental design; (C–F) tumor growth curves, body-weight changes, blood glucose levels, and cumulative food intake in (C) male C57BL/6 mice bearing MC38 colon carcinoma, (D) female C57BL/6 mice bearing MC38 colon carcinoma, (E) male C57BL/6 mice bearing MCA205 sarcoma, and (F) female BALB/c mice bearing EMT6 breast carcinoma. (G) Prophylactic MedDiet intervention in male C57BL/6 mice challenged with MC38 cells: schematic of the experimental design; tumor growth curves, body-weight changes, blood glucose levels, and cumulative food intake. (*P < 0.05; **P < 0.01; ***P < 0.001; ****P < 0.0001)

To further evaluate the influence of sex and tumor type, we tested the MedDiet in female mice bearing either subcutaneous MC38 colon carcinoma (Fig. 1E) or orthotopic EMT6 mammary carcinoma (Fig. 1F). In both models, MedDiet significantly suppressed tumor growth, confirming its efficacy across distinct tumor contexts and biological sexes. The consistency of this response across models underscores a broad-spectrum antitumor effect that is largely independent of host sex, tissue origin, or tumor genetic background.

We next investigated whether MedDiet could confer persistent or preventive antitumor benefits. Remarkably, a 4-week preconditioning regimen of MedDiet prior to tumor implantation, followed by reversion to a standard control diet, was sufficient to maintain durable tumor suppression (Fig. 1G). This outcome occurred without measurable changes in metabolic parameters, implying that transient dietary intervention can reprogram systemic or microenvironmental factors in a way that sustains long-term antitumor protection. Such durability suggests that MedDiet induces a form of nutritional memory, possibly through lasting changes in immune or microbial networks.

Collectively, these results establish that MedDiet exerts a robust, reproducible, and partially preventive antitumor effect across diverse murine models. By decoupling caloric intake from nutrient quality, our study highlights how specific macronutrient compositions—particularly fiber and omega-3–rich lipids—can reshape the metabolic and immune landscape to constrain tumor progression.

### The antitumor effect of MedDiet depends on adaptive immunity

The robust and reproducible tumor-suppressive effects of MedDiet across multiple murine models prompted us to investigate its underlying mechanism of action. Building on prior evidence that dietary patterns can profoundly modulate antitumor immunity by altering immune cell metabolism and inflammatory tone(Sharma et al., 2024; Zhang et al., 2023), we hypothesized that the antitumor efficacy of MedDiet is mediated primarily through adaptive immune mechanisms. To directly test this hypothesis, we implanted subcutaneous MC38 colon carcinoma into immunodeficient nude mice, which lack functional T cells (Fig. 2A). In stark contrast to immunocompetent hosts, MedDiet failed to suppress tumor growth in these mice. This finding establishes that an intact adaptive immune system is indispensable for the tumor-inhibitory effects of MedDiet.

**Figure 2.**
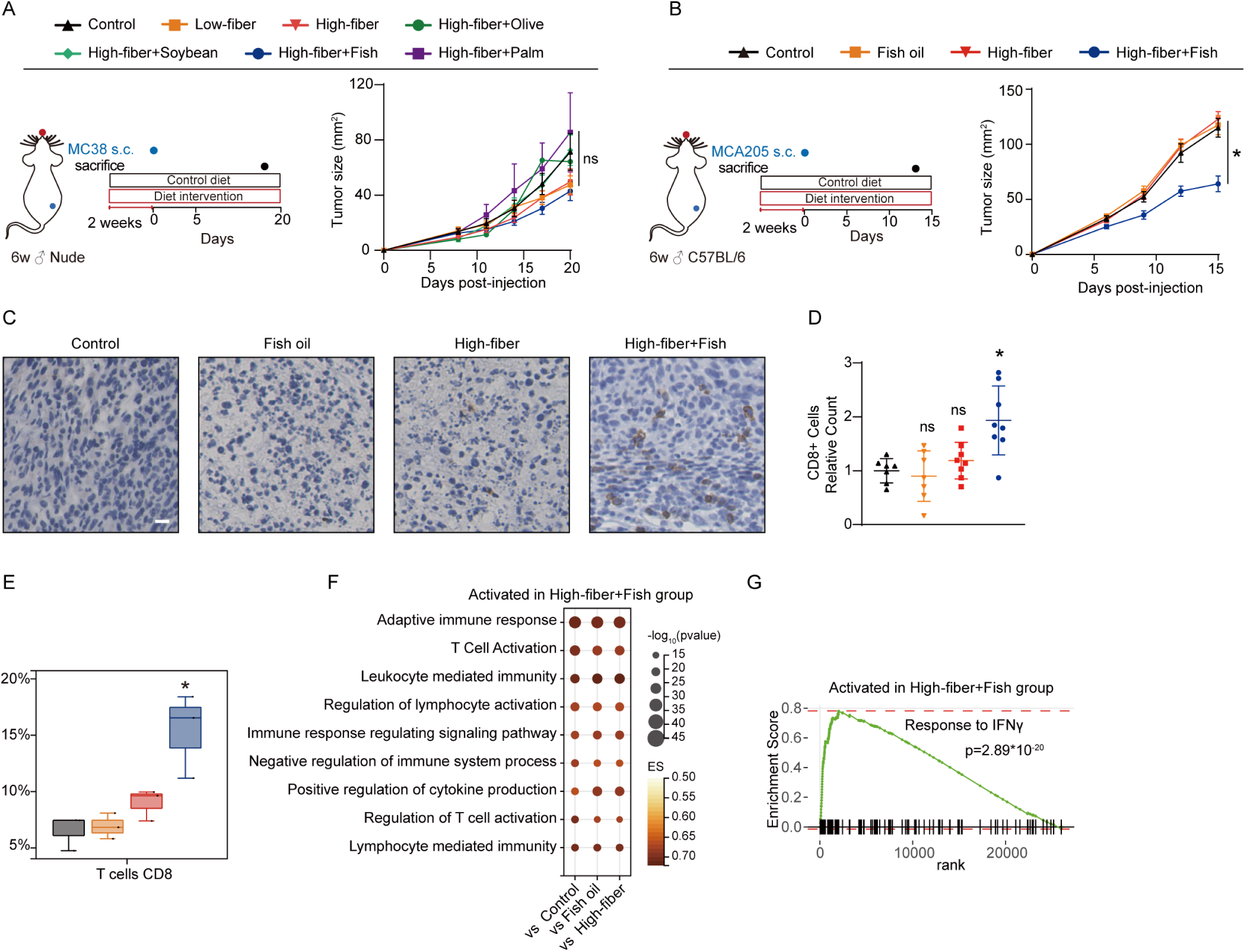
The antitumor effect of the MedDiet depends on adaptive immunity. (A) Nude mice bearing subcutaneous MC38 colon carcinoma subjected to dietary interventions: schematic of the experimental design; tumor growth curves. (B) Male C57BL/6 mice bearing subcutaneous MCA205 fibrosarcoma subjected to dietary interventions: schematic of the experimental design; tumor growth curves. (C, D) (C) Representative images of CD8 immunohistochemistry; (D) quantification of CD8-positive cells. (E) Quantification of CD8^+^ T-cell infiltration levels analyzed using CIBERSORT based on RNA-seq data. (F) Bubble plot displaying pathways upregulated in the MedDiet group compared to other groups, as determined by GSEA. (G) GSEA comparing IFNγ response pathway activation between the MedDiet and control groups. (Scale bar, 20 μm. *P < 0.05; **P < 0.01; ***P < 0.001; ****P < 0.0001)

We next sought to identify which nutritional components of MedDiet contribute to this immunological activation. To this end, we systematically compared the effects of high-fiber and fish oil–enriched diets, administered individually or in combination, in tumor-bearing mice (Fig. 2B). Neither high fiber nor fish oil alone significantly altered tumor growth, suggesting that each nutrient in isolation is insufficient to drive an antitumor response. However, the combination of fiber and fish oil—the defining feature of the MedDiet—recapitulated its full antitumor efficacy, resulting in marked suppression of tumor progression. These findings reveal a synergistic interaction between dietary fiber and omega-3–rich lipids in orchestrating host antitumor responses, likely through metabolic cross-talk between the gut microbiota and the immune system.

To delineate the cellular basis of this response, we examined immune infiltration within the tumor microenvironment (TME). Immunohistochemical analysis revealed a significant increase in CD8⁺ T cell density in tumors from MedDiet-fed mice compared to controls (Fig. 2C, D). Consistent with this observation, bulk transcriptomic profiling of tumor tissues demonstrated selective enrichment of CD8⁺ T cell–associated gene signatures in the MedDiet group, without major shifts in other immune lineages such as macrophages or regulatory T cells (Fig. 2E, Fig. S1). Functional pathway enrichment analysis further highlighted activation of adaptive immune programs, including genes associated with T cell activation, cytotoxicity, and antigen presentation (Fig. 2F). Notably, interferon-γ (IFNγ)–driven signaling pathways were among the most significantly upregulated, consistent with heightened effector function of cytotoxic T lymphocytes (Fig. 2G). Together, these findings demonstrate that the antitumor effects of MedDiet are critically dependent on adaptive immunity, specifically through the mobilization and functional activation of CD8⁺ effector T cells within the TME. The synergy between dietary fiber and fish oil appears to prime an immune-permissive state that enhances cytotoxic T cell infiltration and IFNγ-mediated effector activity, thereby reinforcing host immune surveillance.

### MedDiet-induced enrichment of *B. thetaiotaomicron* drives tumor suppression

The durable and reproducible antitumor activity of MedDiet (Fig. 1G), together with the well-established influence of dietary patterns on gut microbial composition(Mao et al., 2023), led us to hypothesize that its therapeutic effects are mediated, at least in part, through modulation of the gut microbiota. To directly test this possibility, we depleted commensal microbes by pretreating mice with a broad-spectrum antibiotic cocktail, thereby disrupting intestinal microbial communities (Fig. 3A). Remarkably, this intervention abolished the tumor-suppressive effects of MedDiet, as evidenced by the restored growth of MC38 tumors in antibiotic-treated mice. These results demonstrate that an intact gut microbiome is essential for the antitumor efficacy of MedDiet, implicating microbiota-derived signals as critical mediators of its immune-modulatory function.

**Figure 3.**
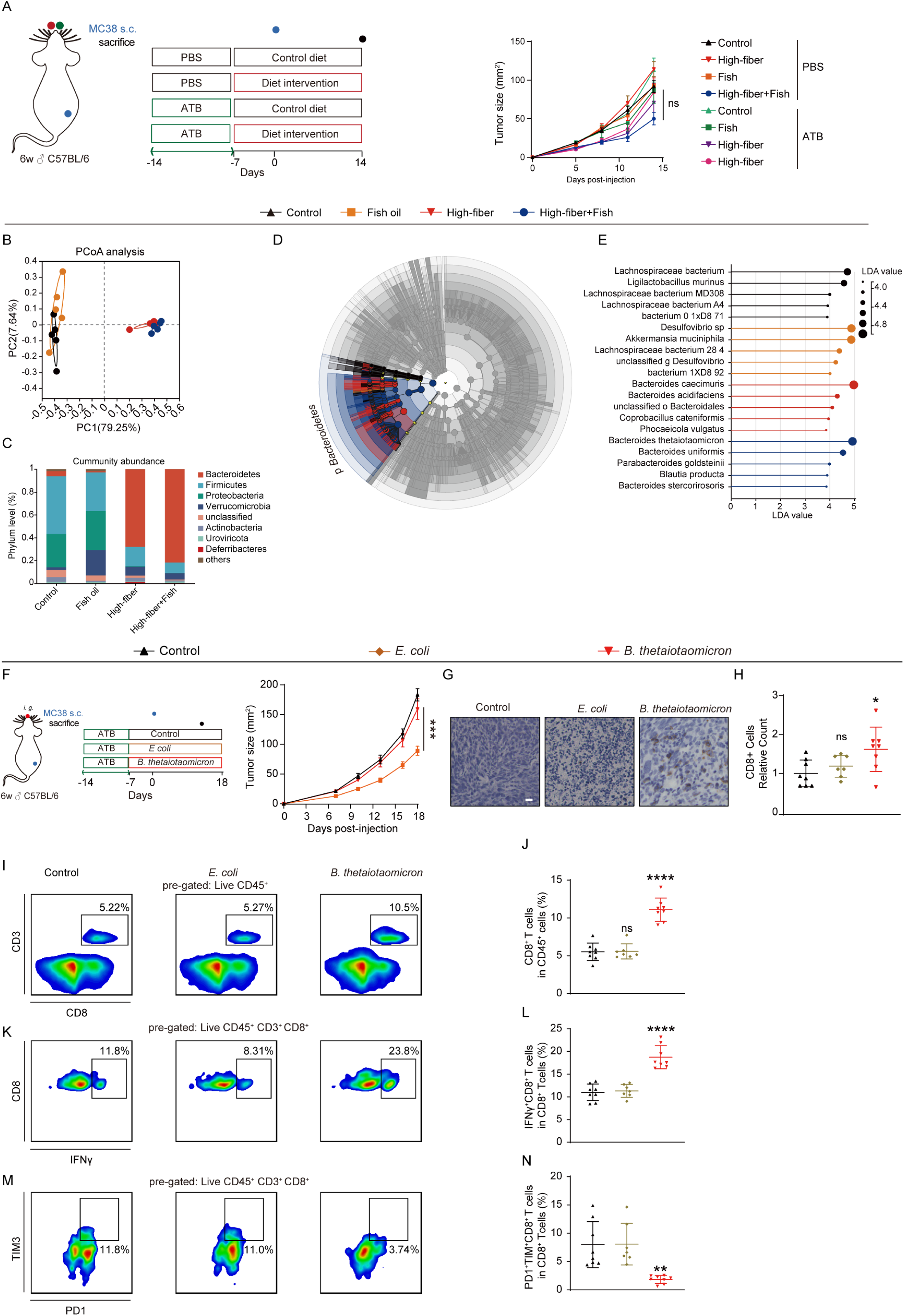
Gut microbiota-dependent mechanisms reveal the pivotal role of *B. thetaiotaomicron* in MedDiet-mediated antitumor effects. (A) Subcutaneous MC38 colon carcinoma model subjected to dietary interventions following antibiotic-mediated depletion of gut microbiota: schematic of the experimental design and tumor growth curves. (B–E) Metagenomic sequencing analysis of gut contents from diet-intervened mice: (B) PCoA; (C) bacterial taxonomic composition at the genus level; (D, E) LEfSe results—(D) LDA score histogram and (E) cladogram. (F) Subcutaneous MC38 colon carcinoma model subjected to bacterial gavage interventions: schematic of the experimental design and tumor growth curves. (G, H) (G) Representative images of CD8 immunohistochemistry; (H) quantification of CD8-positive cells. (I–N) Flow cytometric analysis of intratumoral CD8⁺ T cells: (I, J) assessment of intratumoral CD8⁺ T cells—(I) representative plots and (J) quantification of CD8⁺ T-cell infiltration; (K–N) assessment of intratumoral effector CD8⁺ T cells (IFNγ⁺ CD8⁺ T cells) and exhausted CD8⁺ T cells (PD1⁺ TIM3⁺ CD8⁺ T cells)—(K, M) representative plots, (L) quantification of effector CD8⁺ T cells, (N) quantification of exhausted CD8⁺ T cells. (Scale bar, 20 μm. *P < 0.05; **P < 0.01; ***P < 0.001; ****P < 0.0001)

To identify the specific microbial taxa associated with MedDiet-induced tumor suppression, we performed metagenomic sequencing of fecal samples collected after dietary intervention (Fig. 3B). At the genus level, MedDiet-fed mice exhibited a distinct microbial profile dominated by *Bacteroides* species, in sharp contrast to control diet groups (Fig. 3C). Comparative taxonomic analysis using linear discriminant analysis effect size (LEfSe) revealed a significant enrichment of *Bacteroides* in the MedDiet cohort relative to all other dietary conditions (Fig. 3D, E). Among these, *B. thetaiotaomicron* emerged as the most prominently expanded species—a commensal bacterium known for its robust metabolic versatility, capacity for mucosal adaptation, and previously reported immunomodulatory properties.

To establish a causal relationship between *B. thetaiotaomicron* and tumor suppression, we colonized MC38 tumor-bearing mice with *B. thetaiotaomicron* administration. Strikingly, supplementation with *B. thetaiotaomicron* markedly inhibited tumor growth (Fig. 3F), phenocopying the antitumor effects observed under MedDiet conditions. Immunohistochemistry and flow cytometric analyses revealed a profound increase in intratumoral CD8⁺ T cell infiltration, accompanied by enhanced spatial proximity of T cells to tumor nests (Fig. 3G–J). Moreover, functional profiling of tumor-infiltrating lymphocytes demonstrated a significant rise in IFNγ–producing CD8⁺ effector T cells (Fig. 3K, 3L), together with a reduction in exhausted PD1⁺TIM3⁺ CD8⁺ T cells (Fig. 3M, 3N). These findings indicate that *B. thetaiotaomicron* not only enhances CD8⁺ T cell recruitment but also rejuvenates their functional state within the tumor microenvironment (TME). By contrast, colonization with *Escherichia coli* (*E. coli*), used as a negative control, failed to affect tumor growth, T cell infiltration, or activation status (Fig. 3G–N), confirming the species specificity of this immunostimulatory effect.

Collectively, these results establish that the tumor-suppressive effects of MedDiet are microbiota dependent, with *B. thetaiotaomicron* emerging as a central microbial effector that bridges dietary modulation to immune activation. By reshaping the intestinal ecosystem, MedDiet selectively enriches *B. thetaiotaomicron*, which in turn enhances cytotoxic T cell infiltration, restores effector function, and suppresses tumor progression.

### Tryptophan metabolite 3-IAA links microbiota to immune activation

Microbial metabolites have emerged as pivotal mediators of the crosstalk between diet, immunity, and tumor biology(Trefny et al., 2025). To elucidate whether microbial metabolism underlies the antitumor effects of the MedDiet, we performed untargeted plasma metabolomic profiling of diet-treated mice (Fig. 4A). This analysis revealed a prominent activation of the tryptophan metabolic pathway in the MedDiet group (Fig. 4B), characterized by a marked accumulation of the indole derivative 3-IAA (Fig. 4C). Because 3-IAA is a known *B. thetaiotaomicron*–derived metabolite(Tintelnot et al., 2023), we hypothesized that MedDiet-mediated enrichment of *B. thetaiotaomicron* elevates systemic 3-IAA levels. Indeed, quantitative assays confirmed that oral administration of *B. thetaiotaomicron* markedly increased circulating 3-IAA concentrations in mice (Fig. 4D), establishing a direct metabolic link between dietary microbiota remodeling and systemic metabolite production.

**Figure 4.**
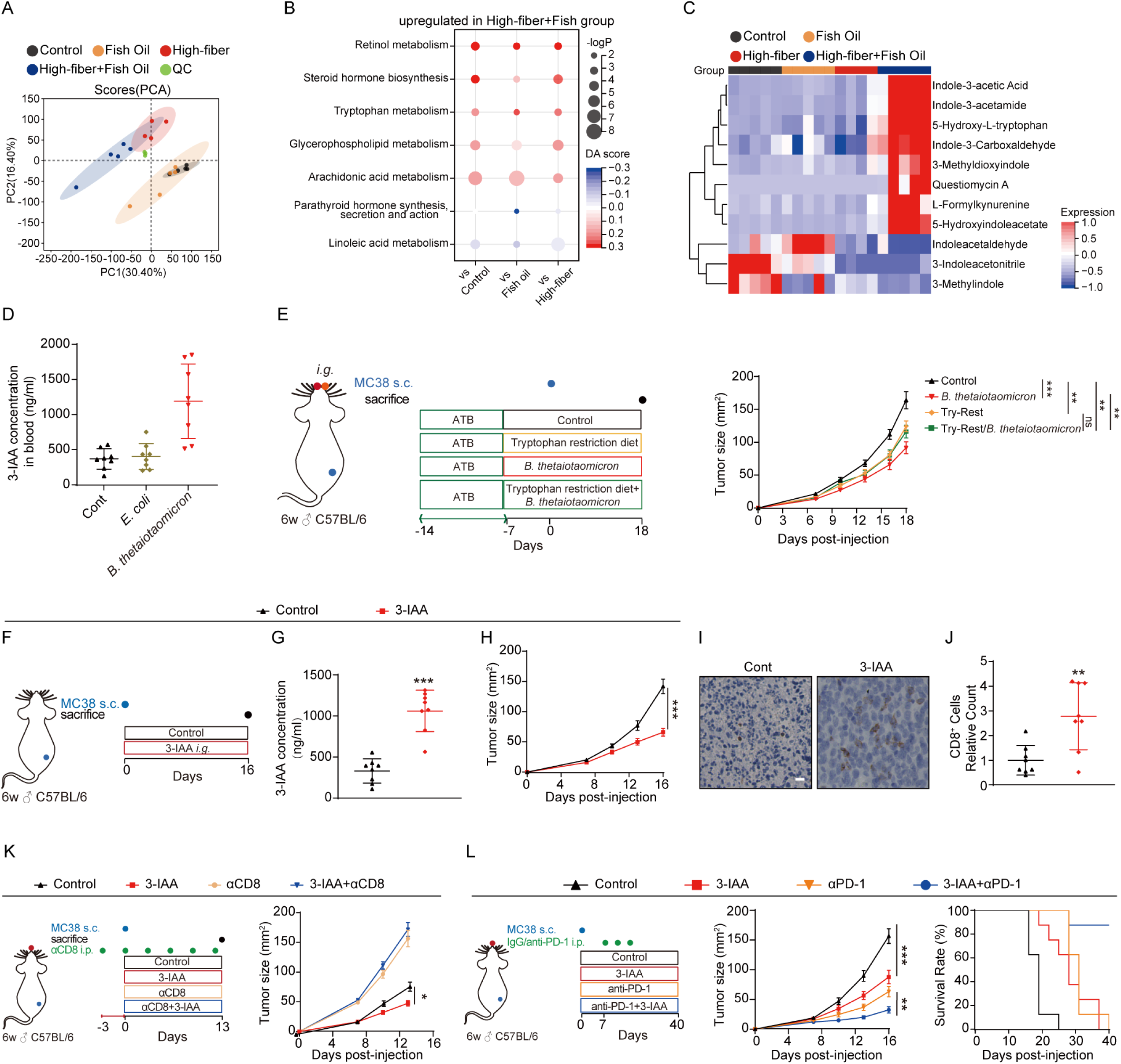
Metabolomics reveal 3-IAA as a key effector molecule in the microbiota–immune axis. (A, B) Untargeted metabolomic analysis of plasma samples from diet-intervened mice: (A) PCA; (B) KEGG enrichment analysis highlighting pathways upregulated in the MedDiet group; (C) heatmap of abundance of differential metabolites in the tryptophan metabolism pathway. (D) Quantification of 3-IAA concentrations in plasma samples from bacteria-gavaged mice using ELISA. (E) Subcutaneous MC38 colon carcinoma model subjected to *B. thetaiotaomicron* gavage and/or tryptophan-restricted diet: schematic of the experimental design; tumor growth curves. (F–J) Subcutaneous MC38 colon carcinoma model treated with oral 3-IAA: (F) schematic of the experimental design; (G) quantification of plasma 3-IAA concentrations using ELISA; (H) tumor growth curves; (I, J) (I) representative images of CD8 immunohistochemistry; (J) quantification of CD8-positive cells. (K) Subcutaneous MC38 colon carcinoma model subjected to CD8 in vivo neutralizing antibody and/or oral 3-IAA treatment: schematic of the experimental design; tumor growth curves. (L) Subcutaneous MC38 colon carcinoma models treated with 3-IAA, anti-PD1 antibody, or their combination: schematic of the treatment regimen; tumor growth curves over time; Kaplan–Meier survival curves. (Scale bar, 20 μm. *P < 0.05; **P < 0.01; ***P < 0.001; ****P < 0.0001)

To probe the functional significance of tryptophan metabolism in tumor control, we next evaluated its contribution in vivo. In MC38 tumor-bearing mice, both *B. thetaiotaomicron* supplementation and tryptophan restriction independently inhibited tumor growth. Strikingly, however, the antitumor efficacy of *B. thetaiotaomicron* was substantially blunted under tryptophan-restricted conditions (Fig. 4E). These results indicate that the bacterium’s tumor-suppressive effects rely on host tryptophan availability, implicating tryptophan-derived metabolites—particularly 3-IAA—as key mediators of this microbiota-dependent immune regulation.

To directly assess the role of 3-IAA, we administered the metabolite orally to tumor-bearing mice (Fig. 4F). Systemic supplementation of 3-IAA significantly increased plasma 3-IAA levels (Fig. 4G) and produced a pronounced inhibition of MC38 tumor growth (Fig. 4H). Immunohistochemical analysis revealed robust CD8⁺ T cell infiltration within the tumor microenvironment of 3-IAA–treated mice (Fig. 4I, 4J), consistent with enhanced adaptive immune activation. Notably, depletion of CD8⁺ T cells completely abolished the antitumor activity of 3-IAA (Fig. 4K), demonstrating that its efficacy depends on cytotoxic T cell–mediated immunity. Given its capacity to amplify CD8⁺ T cell function, we next tested whether 3-IAA could potentiate immune checkpoint blockade (ICB) therapy. In MC38 tumor-bearing mice, combination treatment with 3-IAA and anti–PD-1 antibody synergistically suppressed tumor growth beyond either treatment alone (Fig. 4L). These findings highlight 3-IAA as a functional bridge between microbial metabolism and immune activation.

Together, these results identify 3-IAA as a key effector metabolite within the MedDiet–microbiota–immune axis. By linking B. thetaiotaomicron enrichment to systemic tryptophan metabolism and enhanced CD8⁺ T cell–driven immunity, 3-IAA provides a direct molecular conduit through which dietary and microbial factors converge to suppress tumor progression.

### 3-IAA rejuvenates T cell by alleviating the ISR

To delineate the cellular mechanisms by which the microbiota-derived metabolite 3-IAA mediates its antitumor effects, we first examined its direct impact on tumor and immune cells in vitro. Treatment with 3-IAA did not significantly affect the proliferation of MC38 tumor cells (Fig. S2A) or promote CD8⁺ T cell expansion (Fig. S2B, S2C), suggesting that its efficacy does not stem from direct cytostatic activity or global T cell proliferation. Instead, given our in vivo findings that *B. thetaiotaomicron* supplementation enhanced effector CD8⁺ T cell responses while diminishing exhausted subsets (Fig. 3K–3N), we hypothesized that 3-IAA mitigates T cell exhaustion, a critical barrier to sustained antitumor immunity. To model this process ex vivo, purified CD8⁺ T cells were subjected to either acute stimulation (control) or chronic stimulation (exhaustion) via prolonged anti-CD3 engagement, thereby recapitulating persistent antigenic exposure typical of the tumor microenvironment(Belk et al., 2022) (Fig. 5A). Under acute stimulation, 3-IAA induced only minor reductions in exhaustion marker expression. However, under chronic stimulation, 3-IAA profoundly reversed the exhausted phenotype, increasing the proportion of functional effector CD8⁺ T cells while reducing terminally exhausted subsets (Fig. 5B–5D). To determine whether this phenotypic restoration translated into improved function, we performed co-culture cytotoxicity assays using OT-1 T cells and MC38-OVA tumor cells. While 3-IAA did not enhance killing at a 1:1 effector-to-target ratio, it significantly improved cytotoxicity under high tumor burden conditions that typically induce exhaustion (Fig. 5E). Notably, chronically stimulated T cells treated with 3-IAA exhibited robust cytolytic activity across multiple effector-to-target ratios, confirming that 3-IAA reestablishes T cell effector competence under stress conditions that normally suppress immune function (Fig. 5E).

**Figure 5.**
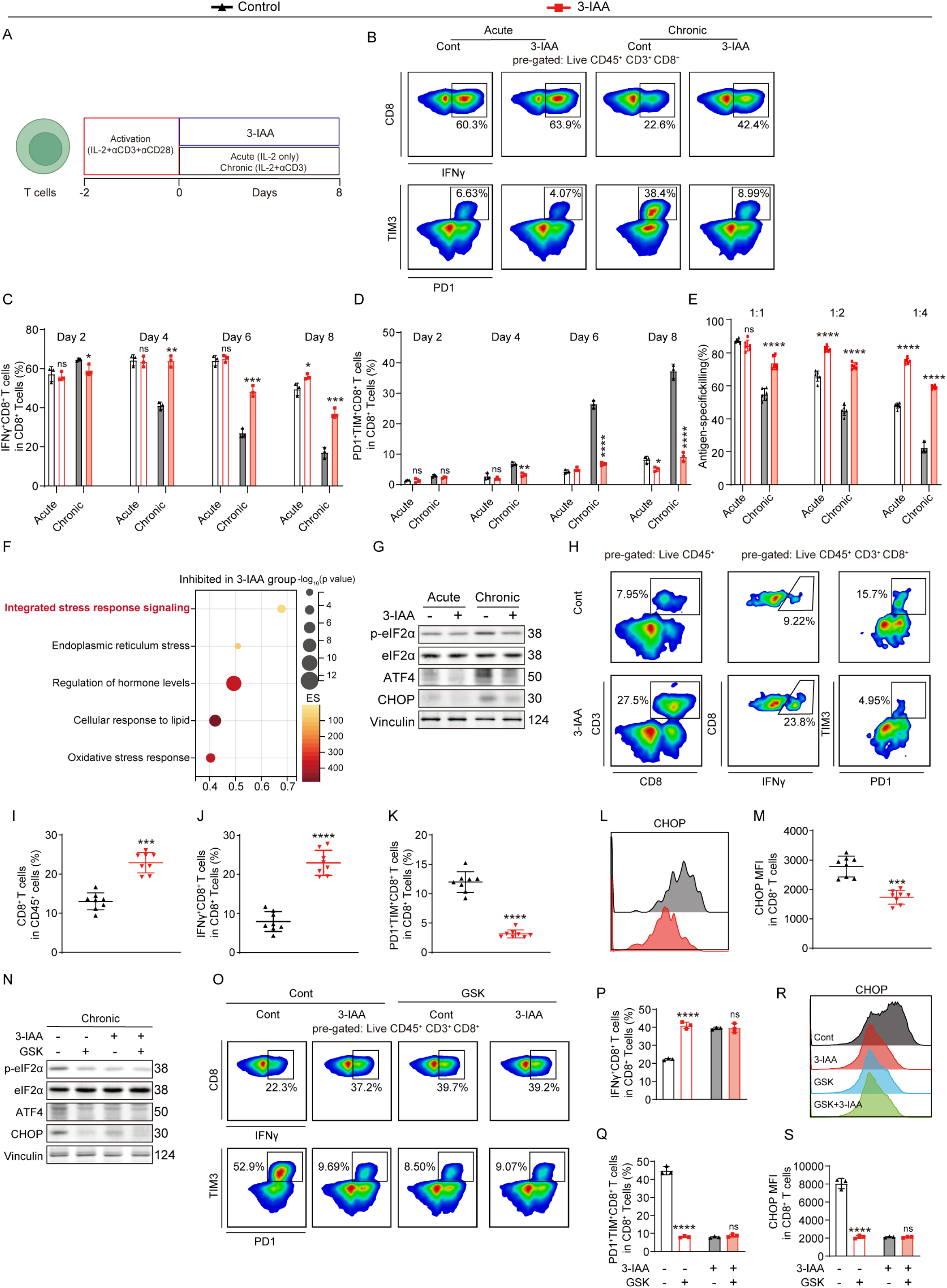
3-IAA alleviates the ISR to inhibit CD8^+^ T cell exhaustion. (A–D) Flow cytometric analysis of CD8⁺ T cells under acute or chronic stimulation over 8 days with or without 3-IAA treatment: (A) schematic of T-cell activation, stimulation, and treatment; (B) representative plots on day 8; (C) quantification of effector CD8⁺ T cells; (D) quantification of exhausted CD8⁺ T cells. (E) Lactate dehydrogenase (LDH) release assay evaluating cytotoxicity of 3-IAA-treated OT1 mouse-derived T cells against MC38-OVA cells. (F) Bubble plot illustrating gene set enrichment analysis (GSEA) of pathways downregulated in chronically stimulated T cells following 3-IAA treatment. (G) Western blot analysis showing expression levels of p-eIF2α, total eIF2α, ATF4, and CHOP in acutely and chronically stimulated T cells with or without 3-IAA treatment. (H–M) Flow cytometric analysis of intratumoral CD8^+^ T cells following oral administration of 3-IAA, evaluating infiltration, effector function, exhaustion levels, and CHOP expression: (H) representative plots; (I) quantification of CD8^+^ T-cell infiltration; (J) quantification of effector CD8^+^ T cells; (K) quantification of exhausted CD8^+^ T cells; (L) histogram of CHOP expression levels; (M) quantification of CHOP expression. (N) Western blot analysis of p-eIF2α, total eIF2α, ATF4, and CHOP in chronically stimulated T cells treated with 3-IAA, GSK, or their combination. (O–S) Flow cytometric analysis of CD8^+^ T cells treated in vitro with 3-IAA, GSK, or their combination, assessing effector function, exhaustion levels, and CHOP expression: (O) representative plots; (P) quantification of effector CD8^+^ T cells; (Q) quantification of exhausted CD8^+^ T cells; (R) histogram of CHOP expression levels; (S) quantification of CHOP expression. (*P < 0.05; **P < 0.01; ***P < 0.001; ****P < 0.0001).

To elucidate the molecular basis of this T cell rejuvenation, we next conducted transcriptomic profiling of chronically stimulated CD8⁺ T cells treated with 3-IAA. RNA sequencing revealed a marked suppression of the ISR pathway in the 3-IAA group (Fig. 5F), suggesting that modulation of cellular stress signaling underlies the restoration of effector function. Western blot analysis confirmed that chronic stimulation activated canonical ISR effectors—including phosphorylated eIF2α, ATF4, and CHOP—whereas 3-IAA treatment normalized their expression to near-baseline levels (Fig. 5G). In vivo experiments corroborated these findings: oral administration of 3-IAA significantly enhanced intratumoral CD8⁺ T cell infiltration (Fig. 5H, I), increased the effector CD8⁺ T cell proportion (Fig. 5H, J), decreased the exhausted subset (Fig. 5H, K), and lowered CHOP expression levels (Fig. 5L, M).

Although the ISR can be activated by four upstream kinases (PERK, GCN2, PKR, and HRI), PERK is reported to be the predominant sensor activated by metabolic and hypoxic stress in tumor-infiltrating CD8⁺ T cells(Cao et al., 2019; Chen et al., 2025; Wang et al., 2025). To further test this mechanism, we pharmacologically inhibited the ISR sensor PERK using GSK2606414 (GSK), a selective PERK inhibitor known to suppress ISR activation(Chen et al., 2025). PERK inhibition recapitulated the effects of 3-IAA, reducing exhaustion and restoring effector T cell identity, while combined treatment with GSK and 3-IAA yielded no additive benefit (Fig. 5N). Flow cytometric analyses further substantiated these observations: both 3-IAA and GSK increased effector CD8⁺ T cell frequencies, reduced exhausted subsets (Fig. 5O–5Q), and markedly decreased CHOP expression (Fig. 5R, 5S). Together, these data confirm that 3-IAA rejuvenates T cell function primarily through suppression of the PERK–ISR axis.

Collectively, our findings reveal a previously unrecognized immunometabolic mechanism by which a microbiota-derived metabolite restores T cell functionality. By mitigating ISR-driven exhaustion, 3-IAA relieves the metabolic and proteostatic constraints imposed by chronic activation, thereby reprogramming exhausted CD8⁺ T cells toward a renewed effector phenotype. This work establishes 3-IAA as a metabolic adjuvant capable of reinvigorating antitumor immunity through targeted modulation of cellular stress responses, bridging microbiota metabolism with host immune resilience.

### A MedDiet score correlates with enhanced antitumor immunity and predicts favorable outcomes in human cancers

To evaluate the association of the MedDiet with human tumorigenesis and therapeutic responses, we further generated a MedDiet Score based on MedDiet-mediated transcriptomic alterations. Subsequently, we interrogated TCGA to determine whether MedDiet Score expression correlates with the tumor immune microenvironment in cancer patients. Pan-cancer analysis revealed that the MedDiet Score exhibited significant positive correlations with the degree of CD8⁺ T cell infiltration across 32 cancer cohorts in TCGA (Fig. 6A). Concurrently, the MedDiet Score showed positive associations with T cell cytotoxicity, central memory, and effector memory phenotypes (Fig. 6B-D). Prognostic analysis indicated that high MedDiet Score expression was associated with favorable outcomes in bladder urothelial carcinoma, esophageal carcinoma, glioblastoma multiforme and low-grade glioma, lung adenocarcinoma, metastatic melanoma, and uveal melanoma (Fig. 6E).

**Figure 6.**
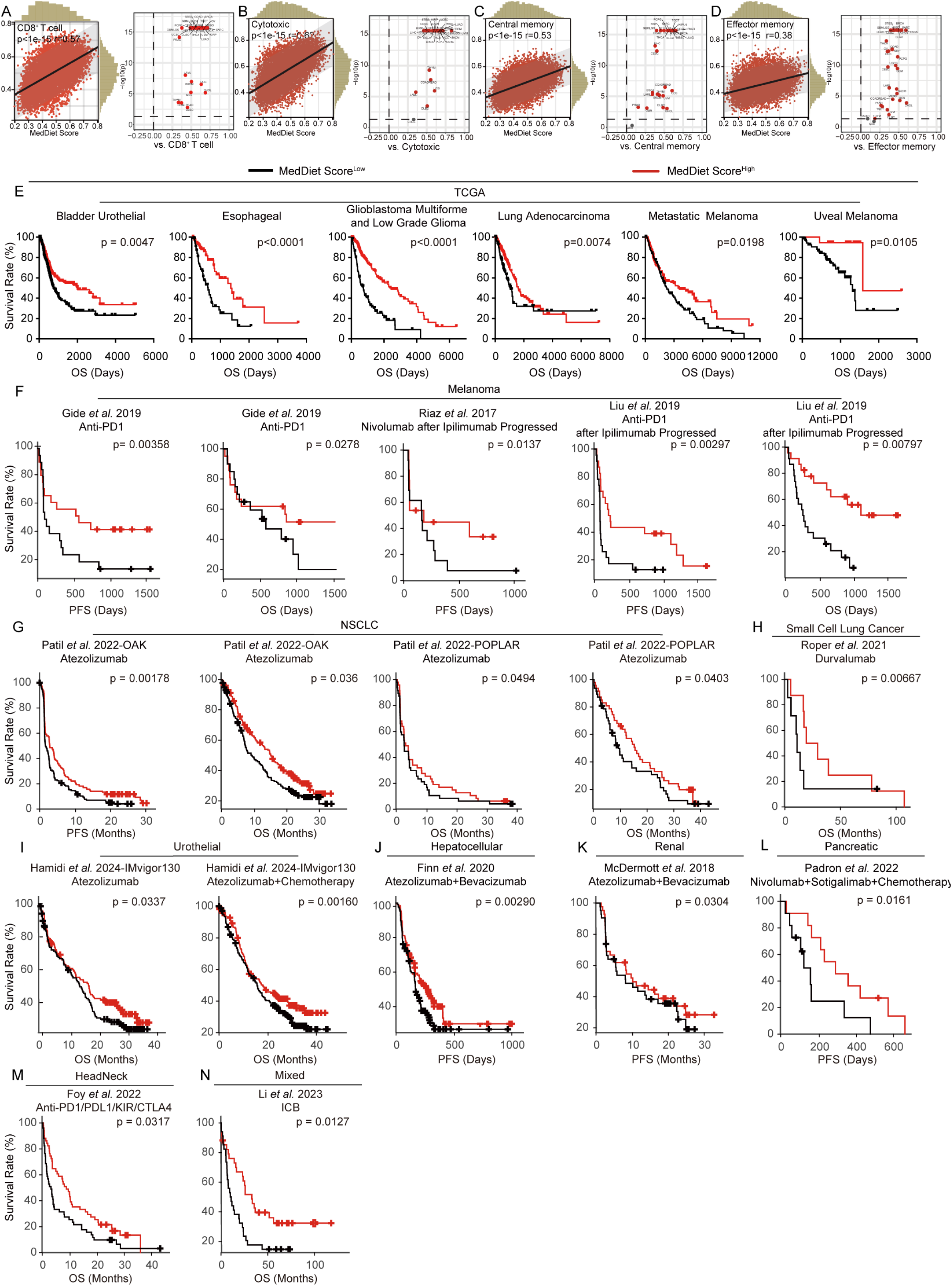
A MedDiet score correlates with enhanced antitumor immunity and predicts favorable outcomes in human cancers. (A–D) Correlation analyses between MedDiet scores and (A) CD8^+^ T-cell infiltration levels, (B) cytotoxic scores, (C) central memory scores, and (D) effector memory scores.in patients from the TCGA pan-cancer cohort (E) Kaplan–Meier survival curves for patients with high versus low MedDiet score-expressing tumors. (F–N) Kaplan–Meier survival curves for patients with high versus low MedDiet score-expressing tumors following immunotherapy: (F) melanoma; (G) non-small cell lung cancer (NSCLC); (H) small cell lung cancer; (I) urothelial carcinoma; (J) hepatocellular carcinoma; (K) renal cell carcinoma; (L) pancreatic cancer; (M) head and neck cancer; and (N) mixed cancer cohorts.

We next assessed the predictive value of the MedDiet Score across multiple public cancer immunotherapy datasets. The results demonstrated that high MedDiet Score expression was indicative of improved post-immunotherapy prognosis in cohorts including melanoma (Fig. 6F), non-small cell lung cancer (Fig. 6G), small cell lung cancer (Fig. 6H), urothelial carcinoma (Fig. 6I), hepatocellular carcinoma (Fig. 6J), renal cell carcinoma (Fig. 6K), pancreatic cancer (Fig. 6L), head and neck cancer (Fig. 6M), and mixed cancer types (Fig. 6N).

Furthermore, we analyzed the association between 3-IAA concentrations and pancreatic cancer prognosis in publicly available cohort data. The findings showed that serum 3-IAA levels were significantly higher in chemotherapy responders than in non-responders (Fig. S3A). Moreover, in two independent cohorts, elevated serum 3-IAA concentrations were associated with favorable prognosis in cancer patients following chemotherapy (Fig. S3B–D).

In summary, the MedDiet Score not only reflects the “hotness” of the tumor immune microenvironment but also serves as a reliable pan-cancer, cross-therapeutic prognostic and immunotherapy response predictor, providing novel molecular insights for clinical precision nutritional interventions combined with immunotherapy. Elevated serum 3-IAA levels also associate with favorable chemotherapy outcomes in pancreatic cancer across cohorts.

## Discussion

This study delineates a diet–microbiota–immune axis through which a high-fiber, fish oil–based regimen (MedDiet) elicits durable tumor suppression across multiple murine cancer models. Mechanistically, MedDiet selectively enriches *B. thetaiotaomicron*, which activates the tryptophan metabolic pathway to generate the microbial metabolite 3-IAA. Acting as a key effector molecule, 3-IAA alleviates CD8⁺ T cell exhaustion by suppressing the ISR signaling cascade, thereby restoring cytotoxic activity and enhancing responsiveness to PD-1 blockade (Fig. 7). Collectively, these findings illuminate how rationally designed diets can reprogram immune metabolism and potentiate immunotherapeutic efficacy.

**Figure 7.**
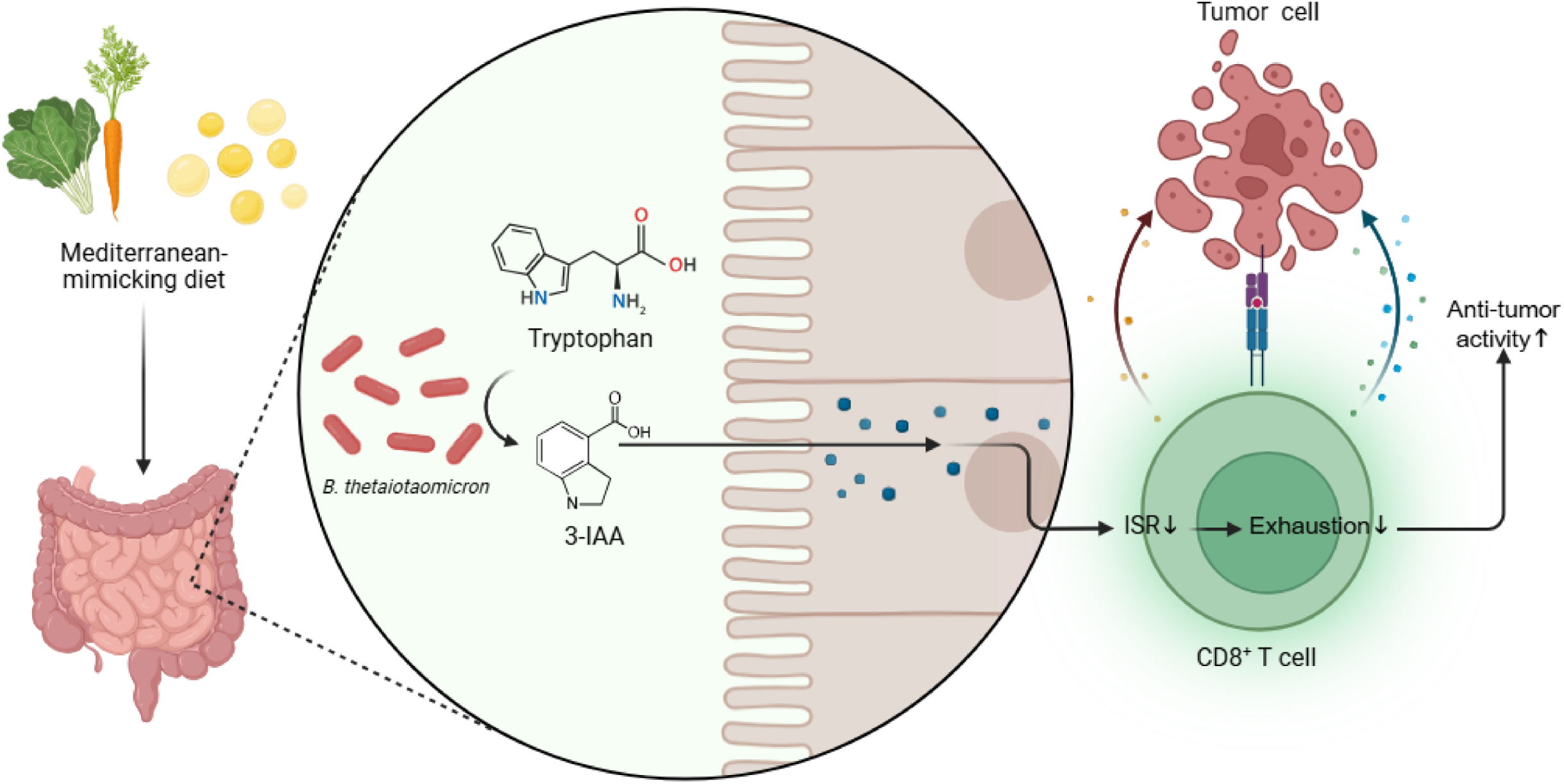
Graphical abstract. This schematic illustrates how a MedDiet suppresses tumor growth by activating a microbiota–metabolite–immune axis centered on gut microbiota-derived 3-IAA, thereby rejuvenating antitumor CD8⁺ T-cell immunity. A Mediterranean-mimicking diet selectively enriches *B. thetaiotaomicron* in the gut microbiota; administration of this bacterium alone recapitulates the diet’s tumor-suppressive effects. Both the MedDiet and B. thetaiotaomicron colonization significantly elevate systemic levels of the tryptophan-derived metabolite 3-IAA. Mechanistically, 3-IAA potentiates CD8⁺ T-cell cytotoxicity and reverses T-cell exhaustion by suppressing the integrated stress response, thereby sustaining robust antitumor immunity in vivo.

Our work extends a growing body of research demonstrating that specific dietary interventions can modulate systemic metabolism and antitumor immunity. Distinct nutritional regimens—ranging from high-fiber diets and caloric restriction to ketogenic and low-glycemic approaches—have been shown to influence tumor progression through diverse mechanisms, including altered lipid utilization, insulin and IGF signaling, and modulation of host immunity(Blaževitš et al., 2023; Evan C. Lien et al., 2021; Qin et al., 2024). For example, high dietary fiber intake correlates with improved progression-free survival in patients receiving ICB, particularly in those not consuming probiotics(Spencer et al., 2021). However, the benefits of fermentable fiber are not universal: soluble fibers such as inulin can induce microbiota-dependent hepatocellular carcinoma in dysbiotic mice, underscoring the need for caution when enriching diets with fermentable substrates(Singh et al., 2018; Yang et al., 2024). Conversely, microbiota modulation through non-fermentable fibers or balanced high-fiber diets has been shown to enhance antitumor immunity via activation of the intratumoral IFN-I–NK cell–dendritic cell axis, improving responsiveness to ICB(Lam et al., 2021). Similarly, ketogenic and high-fat dietary paradigms exert tumor-suppressive effects contingent upon lipid composition, as the ratio of saturated to monounsaturated fatty acids critically determines stearoyl-CoA desaturase (SCD) activity and tumor lipid homeostasis(E. C. Lien et al., 2021). Clinical and preclinical studies further reveal that omega-3 fatty acid–rich diets reduce tumor proliferation markers and reprogram macrophages toward a less protumoral phenotype through reactive oxygen species–mediated cytotoxicity(Simpson et al., 2022) (Aronson et al., 2024; Liu et al., 2020). In this context, our findings highlight the synergistic power of combining dietary fiber with fish oil, which not only reshapes the microbiota but also promotes the generation of immunoregulatory metabolites that sustain T cell function. Together, these studies underscore that dietary composition governs tumor growth through a continuum of mechanisms spanning nutrient signaling, metabolic remodeling, and immune reprogramming.

The MedDiet exerts potent antitumor effects in part through remodeling of the gut microbiota, with *B. thetaiotaomicron* emerging as a key mediator linking diet to immune activation. This dominant gut symbiont possesses an extensive repertoire of carbohydrate-active enzymes(Thompson et al., 2015) and tryptophan-catabolic pathways(Russell et al., 2013), enabling the generation of immunoregulatory metabolites such as short-chain fatty acids(Mirretta Barone et al., 2024), sphingolipids(Brown et al., 2025, 2019) and indole derivatives(Sinha et al., 2024). Diet composition profoundly shapes its abundance and function: fermentable fibers and omega-3 polyunsaturated fatty acids promote its expansion(Deehan et al., 2024; Lai et al., 2023; Takeuchi et al., 2025; Watson et al., 2018), whereas sugar-rich diets induce its depletion(Gal-Mandelbaum et al., 2025). These dietary inputs foster a gut ecosystem conducive to anti-inflammatory and antitumor immunity through metabolite production, antigen presentation, and vesicle-mediated immune modulation. Mechanistically, *B. thetaiotaomicron* exerts multifaceted effects on host physiology through multiple routes—by degrading complex glycans via polysaccharide utilization loci (PULs)(Ndeh et al., 2025), producing sphingolipids that modulate intestinal inflammation(Brown et al., 2019; Mirretta Barone et al., 2024), and releasing outer membrane vesicles engage pattern recognition receptors on dendritic and epithelial cells(Gul et al., 2022; Hickey et al., 2015) to drive IL-10–dependent tolerance(Brown et al., 2025) and MHC class II expression(Yang et al., 2025). Notably, T cell responses specific to *B. thetaiotaomicron* have been associated with enhanced efficacy of CTLA-4 blockade, underscoring its role in immunotherapy responsiveness(Vétizou et al., 2015). In addition, single microbial antigens, such as the nutrient-regulated BT4295 protein, can drive regulatory T cell (Treg) differentiation and limit effector T cell activation, providing an additional layer of immune regulation(Wegorzewska et al., 2019). Beyond these immune-modulatory actions, *B. thetaiotaomicron*–derived metabolites, including propionate and 3-IAA, influence histone methylation, redox balance, and CD8⁺ T cell priming(Bhattarai et al., 2018; Ryu et al., 2022; Sinha et al., 2024), thereby bridging microbial metabolism and host transcriptional and immune programs. Under MedDiet conditions, *B. thetaiotaomicron* preferentially channels tryptophan metabolism toward the 3-IAA production. This microbiota–immune interface highlights how defined dietary interventions selectively recalibrate microbial metabolism to enhance host immunosurveillance, offering a conceptual framework for developing microbiota-derived metabolic adjuvants to enhance immunotherapy efficacy.

Microbiota-derived indole metabolites have emerged as pivotal regulators of host immunity and metabolic homeostasis. Among them, 3-IAA represents a particularly potent effector with broad immunometabolic activity(Bala et al., 2024; M et al., 2024). Beyond its established roles in preserving gut barrier integrity and mitigating oxidative stress(Gomes and Scortecci, 2021; Kim et al., 2017), 3-IAA functions in mammalian cells as a radical scavenger that protects against redox imbalance and cellular injury(Arnao et al., 1996; Kim et al., 2017).

Recent evidence links 3-IAA to improved therapeutic response in pancreatic cancer, where elevated systemic levels correlate with enhanced chemotherapy sensitivity. In preclinical models, fecal microbiota transplantation, dietary tryptophan modulation, or direct 3-IAA supplementation similarly potentiate treatment efficacy(Tintelnot et al., 2023), positioning 3-IAA as a microbiota-derived determinant of treatment response. Our findings extend this conceptual framework by identifying CD8⁺ T cells as direct functional targets of 3-IAA within the tumor microenvironment. Chronic antigenic stimulation induces a proteotoxic stress response (PSR) in exhausted T cells, characterized by excessive translation, protein misfolding, and heightened autophagy(Wang et al., 2025). 3-IAA reverses T cell exhaustion by suppressing the ISR. By normalizing ISR markers—including phosphorylated eIF2α, ATF4, and CHOP—3-IAA restores interferon-γ production, degranulation capacity, and cytotoxic function. Importantly, 3-IAA synergizes with PD-1 blockade, amplifying adaptive immunity beyond checkpoint inhibition alone. Together, these findings establish 3-IAA as a pivotal microbiota-derived metabolite that links intestinal metabolism to immune rejuvenation. By mitigating ISR-driven exhaustion, 3-IAA reprograms dysfunctional T cells toward sustained effector competence, offering a conceptual framework in which microbial metabolites can be rationally leveraged as metabolic adjuvants to enhance antitumor immunity.

Despite these advances, several key questions remain open. The molecular sensor or receptor for 3-IAA in mammalian T cells has yet to be identified, warranting structural and signaling studies to map its downstream effectors. The durability, reversibility, and safety of MedDiet-induced microbiota reprogramming in humans also require systematic clinical investigation. Moreover, while our work focuses primarily on CD8⁺ T cells, the tumor microenvironment encompasses diverse immune and stromal populations whose responses to dietary modulation remain incompletely understood. Future efforts should examine how defined nutritional interventions influence regulatory T cells, myeloid populations, and cytokine networks to orchestrate a coordinated immune response. Ultimately, delineating these diet–microbiota–immune interactions may enable the rational design of precision nutritional strategies that synergize with immunotherapy to achieve durable and systemic tumor control.

In summary, our study identifies a defined diet–microbiota–immune circuit through which the Mediterranean diet reprograms host metabolism to sustain cytotoxic T cell function and potentiate immunotherapy. By linking *B. thetaiotaomicron*–derived 3-IAA to alleviation of stress-induced T cell exhaustion, these findings reveal a metabolite-mediated mechanism of immune rejuvenation. Beyond advancing our understanding of nutritional immunology, this work highlights the therapeutic potential of rationally engineered diets and microbial consortia as metabolic adjuvants to augment checkpoint blockade, offering a conceptual framework for precision dietary interventions in cancer treatment.

## Materials and methods

### Cell lines and cell culture

The mouse tumor cell lines MC38, MCA205, and EMT6 were obtained from Professor Guido Kroemer’s laboratory. The MC38-OVA-luc cell line (Cat. No. NM-S13-TM56) was obtained from Shanghai Model Organisms Center, Inc. All cell lines were cultured in DMEM supplemented with 10% (v/v) FBS at 37°C with 5% CO₂.

### Bacterial culture

*E. coli* (strain No. ATCC 25922) and *B. thetaiotaomicron* (strain No. ATCC 29148) was obtained from the American Type Culture Collection (ATCC). *E. coli* was cultured aerobically in LB broth. Frozen *E. coli* stocks were inoculated into 5 mL of LB medium and incubated at 37°C with shaking at 180 rpm. *B. thetaiotaomicron* was cultured anaerobically in brain heart infusion (BHI) broth. Frozen *B. thetaiotaomicron* stocks were inoculated into BHI medium and incubated at 37°C in an anaerobic chamber with a gas mixture of 5% H₂, 10% CO₂, and 85% N₂.

### Mouse models

All animal experiments were approved by the Animal Ethics Committee of Renmin Hospital of Wuhan University, and were performed in accordance with the relevant guidelines. Seven-week-old female wild-type C57BL/6 mice were purchased from the Animal Center of Renmin Hospital of Wuhan University, and OT1 mice (Strain No. T059617) were obtained from GemPharmatech Co., Ltd. Mice were housed in SPF conditions at the Animal Center, Renmin Hospital of Wuhan University, with a 12-hour light/dark cycle at 24°C, and had ad libitum access to sterilized food and water. The custom mouse diet was purchased from Jiangsu XieTong Biological Engineering Co., Ltd. For tumor transplantation, 1 × 10⁶ cells in 100 µL PBS were injected subcutaneously into the right flank of each mouse.

Dietary interventions were initiated 7 days prior to tumor formation and continued until euthanasia. Mice were divided into various groups, including control, high-fiber, low-fiber, and high-fiber combined with different oils (olive oil, soybean oil, fish oil, or palm oil). In the dietary pre-treatment model, mice were fed the designated diets for 28 days prior to tumor cell injection and then switched to a control diet.

Antibiotic treatment was applied via drinking water containing ampicillin (1 g/L, Sigma-Aldrich), vancomycin (0.5 g/L, Sigma-Aldrich), neomycin (1 g/L, Sigma-Aldrich), and metronidazole (1 g/L, Sigma-Aldrich) to deplete gut microbiota. For microbiota supplementation, *B. thetaiotaomicron* and *E. coli* (negative control) were administered by oral gavage starting 7 days prior to tumor inoculation. Mice received 5 × 10^8^ CFU of bacteria suspended in sterile PBS every 2 days.

Tryptophan-restricted diets (0% tryptophan) or control diets (0.2% tryptophan) were provided from 7 days before tumor formation. 3-IAA (500 mg/kg, Sigma-Aldrich) was administered via oral gavage starting on the day of tumor inoculation, with the control group receiving equivalent PBS. CD8⁺ T cells were depleted by intraperitoneal injection of 200 µg anti-CD8 antibody (clone 53-6.7, BioXcell) on days -3, 0, and every 3 days thereafter. The control group received the equivalent isotype control IgG (Clone 2A3, BioXcell).

PD1 blockade treatment was administered when tumor volume reached approximately 20 mm² (day 7 after inoculation), with 200 µg anti-PD1 antibody (clone F.1A12, BioXcell) every 3 days, while the control group received isotype control IgG.

Tumor growth was monitored by measuring the longest (L) and shortest (W) tumor diameters. Tumor area was calculated as (L × W × 4/π).

### Western Blot

Cells were lysed on ice for 30 min using RIPA lysis buffer, followed by centrifugation at 12,000 g for 15 min at 4°C. The supernatant containing soluble proteins was collected for further analysis. Protein samples were mixed with 4× loading buffer and denatured at 100°C for 10 min, then immediately placed on ice to prevent protein aggregation or degradation before electrophoresis. Equal amounts of protein (20-40 μg per lane) were separated using SDS-PAGE and transferred onto 0.2 μm PVDF membranes, which were pre-activated by methanol immersion and rinsed with deionized water. Membranes were blocked with 5% non-fat milk in TBST at room temperature for 1 h with gentle shaking, followed by overnight incubation at 4°C with primary antibodies diluted in 5% non-fat milk TBST. After three 10-min washes with TBST, membranes were incubated with secondary antibodies at room temperature for 1 h and washed again three times. Chemiluminescence detection was performed using an ECL kit (GE Healthcare), and signal acquisition was conducted with a LI-COR imaging system. The primary antibodies used are listed in Table S1.

### Flow cytometry

Tumor tissues were harvested from euthanized mice, weighed, and immediately placed in pre-cooled RPMI-1640 medium on ice. Tumors were mechanically disrupted into 1-2 mm³ fragments using sterile scissors, and a DNase I/Collagenase IV mixture was added for digestion at 37°C with shaking for 1 hour. After digestion, an equal volume of RPMI-1640 containing 10% FBS was added to stop the reaction, and the mixture was filtered through a 70 μm cell strainer to remove undigested tissue. The cell suspension was washed twice with PBS and adjusted to 250 mg/mL in PBS based on the initial tumor weight for further analysis. For intracellular cytokine detection, cells were restimulated in culture medium containing Brefeldin A (5 μg/mL, Biolegend) for 5 hours at 37°C, 5% CO₂ to inhibit cytokine secretion and promote intracellular accumulation. Cell viability was assessed by staining with LIVE/DEAD® Fixable UV Dye (1:1000, Biolegend), followed by 20 min incubation and centrifugation. Fc receptors were blocked using anti-mouse CD16/CD32 monoclonal antibody (1:100, Biolegend) for 15 min at 4°C, followed by surface staining with fluorescently labeled antibodies for 30 min at 4°C in the dark, and the cells were washed by centrifugation. For intracellular staining, cells were fixed and permeabilized using Fix/Perm buffer (Biolegend) for 1 hour at room temperature in the dark, then stained with fluorescently conjugated antibodies against intracellular proteins for 30 min, followed by washing. Flow cytometry was performed using a BD FACS Diva system, and data were analyzed with FlowJo software after filtering samples through a 40 μm mesh to exclude debris and aggregates. The fluorescent antibodies used are listed in Table S2.

### RNA-seq and transcriptome analysis

RNA was extracted from tissue and cell samples using TRIzol reagent. For tissue samples, fresh tissues were snap-frozen in liquid nitrogen, ground into powder using a pre-cooled mortar, and mixed with TRIzol reagent. After vortexing, the samples were incubated at room temperature for 5 min, then homogenized. Following the addition of chloroform, the samples were vigorously shaken and incubated for 3 min at room temperature, then centrifuged at 12,000 rpm for 15 min at 4°C. The upper aqueous phase was transferred to a new tube, and an equal volume of ice-cold isopropanol was added. The mixture was incubated at -20°C for 20-30 min to precipitate RNA, followed by centrifugation at 12,000 rpm for 10 min at 4°C. The RNA pellet was washed with 75% ethanol and resuspended in DEPC-treated water or 0.5% SDS. For cell samples, cells were pelleted, resuspended in TRIzol, and processed as described above for tissue samples. cDNA libraries were constructed using the TruSeq RNA Sample Preparation Kit, starting with 200 ng of RNA. RNA was fragmented, reverse-transcribed using oligo(dT) primers, and second-strand cDNA was synthesized. The libraries were end-repaired, A-tailed, and ligated with sequencing adapters, followed by PCR amplification for enrichment. Library quality was verified using low-range agarose gel electrophoresis to select fragments of the desired size. Cluster generation was performed on the Illumina cBot using the TruSeq PE Cluster Kit v3-cBot-HS, followed by bridge PCR to amplify the cDNA libraries into dense clusters on the flow cell. High-throughput sequencing was performed on the Illumina MiSeq system using the TruSeq SBS Kit v3-HS, with paired-end sequencing to obtain large quantities of sequence data. RNA-seq data analysis was performed using featureCounts to summarize gene expression levels. Gene set enrichment analysis (GSEA) was conducted using the R package ClusterProfiler(Yu et al., 2012), and immune cell infiltration was estimated with CIBERSORT(Chen et al., 2018) based on mRNA expression levels.

### Plasma untargeted metabolomics analysis

Blood samples were collected from the orbital venous plexus of mice under sterile conditions. The blood was immediately transferred to a centrifuge tube containing heparin as an anticoagulant, gently inverted to mix, and stored on ice. After collection, the orbital area was pressed with sterile gauze to stop bleeding. The samples were then centrifuged at 3000 × g for 10 minutes at 4°C, and the plasma was carefully separated, avoiding contamination with red blood cells. Plasma samples were stored at −80°C until analysis.

For plasma sample preparation, samples were thawed slowly on ice and centrifuged at 3000 × g for 10 minutes at 4°C to remove impurities. The supernatant was transferred to a fresh tube and stored at −80°C until further analysis. Metabolite analysis was performed using an UHPLC-Q Exactive system. Mobile phase A consisted of 95% ultrapure water and 5% acetonitrile, both containing 0.1% formic acid; mobile phase B consisted of 47.5% acetonitrile, 47.5% isopropanol, and 5% ultrapure water, also with 0.1% formic acid. The gradient elution program for positive ion mode was as follows: 0–3 minutes, 0–20% B; 3–4.5 minutes, 20–35% B; 4.5–5 minutes, 35–100% B; 5–6.3 minutes, 100% B; 6.3–6.4 minutes, 100–0% B; 6.4–8 minutes, 0% B. For negative ion mode, the program was: 0–1.5 minutes, 0–5% B; 1.5–2 minutes, 5–10% B; 2–4.5 minutes, 10–30% B; 4.5–5 minutes, 30–100% B; 5–6.3 minutes, 100% B; 6.3–6.4 minutes, 100–0% B; 6.4–8 minutes, 0% B. The flow rate was maintained at 0.40 mL/min to ensure optimal separation and detection. Mass spectrometry data were acquired using the Orbitrap detector of the UHPLC-Q Exactive system, with appropriate resolution settings for both positive and negative ion modes to cover a broad range of metabolites. Data were processed by matching accurate mass and fragment ion information against the Human Metabolome Database (HMDB, http://www.hmdb.ca/) and Metlin database (https://metlin.scripps.edu/) for metabolite identification. Metabolic pathway enrichment and network topology analyses were performed based on the KEGG database to interpret the biological significance of the identified metabolites.

### Fecal metagenomic sequencing analysis

Fresh fecal samples were collected using sterile containers and immediately stored at –80 °C to preserve DNA integrity. Genomic DNA was extracted using the PF Mag-Bind Fecal DNA Extraction Kit. Briefly, frozen samples were homogenized with the provided homogenization buffer using a bead mill (6.0 m/s, 30–40 s, 2–3 cycles, with 1 min cooling on ice between cycles) to prevent DNA degradation. Subsequently, lysis buffer and proteinase K were added, and the mixture was incubated at 70 °C for 10–15 min to disrupt cellular structures. After adding the inhibitor removal buffer, samples were centrifuged at 12,000 × g for 5 min to pellet debris. The supernatant was transferred to magnetic bead binding columns, and DNA was bound by adding binding buffer, mixing, and incubating for 5 min. Magnetic separation was used to remove the supernatant, and washing steps were performed using the provided wash buffers to eliminate contaminants. Finally, DNA was eluted with preheated (65 °C) elution buffer, and its concentration and purity were assessed using a NanoDrop spectrophotometer, aiming for A260/A280 ratios between 1.8 and 2.0 and A260/A230 ratios above 1.8. DNA was fragmented to an average size of 400 bp using a sonicator, optimizing power, time, and cycles for uniform distribution. Libraries were constructed using the NEXTFLEX Rapid DNA-Seq Library Prep Kit, following the manufacturer’s protocol, including end repair, A-tailing, adapter ligation, and PCR amplification to ensure quality and compatibility. Library quality was assessed via agarose gel electrophoresis and a bioanalyzer, confirming appropriate size distribution, concentration, and purity. Libraries were then subjected to paired-end sequencing on an Illumina Novaseq 6000 platform, with parameters set to obtain high-quality data. Raw sequencing data were uploaded to the Majorbio Cloud Platform (https://www.majorbio.com) for bioinformatics analysis, which included quality control (removal of reads < 50 bp, quality < 20, or containing N bases), de novo assembly to obtain contiguous genomic sequences, host genome contamination removal through alignment with host databases, and taxonomic composition analysis using database comparisons and classification algorithms to determine microbial species and their relative abundances.

### ELISA

Mouse plasma samples were centrifuged at 3,000 × g for 10 min at 4 °C to remove impurities; the supernatant was transferred to clean tubes and kept on ice. The 3-IAA ELISA Kit (Shanghai Win Win Biotechnology Co., Ltd) was equilibrated to room temperature, and standard solutions were prepared according to the manufacturer’s instructions. Standard samples or plasma were added to pre-coated 96-well plates, with wells designated for blanks, standards, and samples, each in triplicate. The plate was incubated at 37 °C for 30 min to allow antigen-antibody binding. After incubation, the liquid was discarded, and the plate was washed five times with washing buffer, tapping gently on absorbent paper after each wash to remove unbound substances. Biotin-labeled detection antibody was added to each well, ensuring complete coverage, and the plate was incubated at 37 °C for 30 min. Following incubation, the plate was washed five times as before. Color development was initiated by adding substrate solution to each well, incubating in the dark at 37 °C for 10 min, and monitoring color change to ensure sufficient reaction without overdevelopment. The reaction was terminated by adding 50 μL of stop solution, turning the color from blue to yellow. Absorbance (OD) was measured using a microplate reader, recording values for standard and sample wells. A standard curve was constructed by plotting standard concentrations against corresponding OD values, and sample 3-IAA concentrations were determined through linear regression analysis.

### CCK8

Cells were seeded in 96-well plates at a density of 5,000 cells per well, adding 200 μL of culture medium to each well to ensure uniform distribution and adherence. To minimize intra-group variability, each sample was prepared in six replicates. An equal volume of PBS was added to the outer wells to prevent evaporation-induced alterations in medium composition and cellular microenvironment. 3-IAA solution was added to the appropriate wells to achieve a final concentration of 1 μg/mL, and cells were incubated for a specified duration as determined by the experimental design. Following incubation, the culture medium was removed, and 100 μL of CCK-8 reagent (Beyotime) was added to each well. The plate was then incubated at 37 °C in the dark for 1 hours to allow for the formation of the formazan product by mitochondrial dehydrogenases in viable cells. Absorbance at 450 nm was measured using a microplate reader, and the optical density values were recorded to assess cell viability and proliferation.

### T cell isolation and activation

Spleen tissues were isolated from experimental mice under sterile conditions and placed in a 5 mL petri dish containing pre-chilled PBS. The spleens were minced using sterile scissors and forceps, followed by mechanical dissociation using a pestle or a 70 μm cell strainer to prepare a single-cell suspension. The suspension was transferred to a 15 mL centrifuge tube and centrifuged at 300 × g for 5 minutes at room temperature to discard the supernatant. The cell pellet was resuspended in 5 mL of red blood cell lysis buffer, incubated at room temperature for 3-5 minutes, and gently mixed. After terminating the lysis by adding an equal volume of PBS, the cells were centrifuged at 300 × g for 5 minutes. The washing step with PBS was repeated to remove residual red blood cell debris. If incomplete red blood cell removal occurred, the lysis step could be extended up to 7 minutes to avoid excessive T cell damage. Finally, the cells were resuspended in pre-chilled complete medium. The T cell culture medium was prepared by mixing 100 mL FBS, 0.11 g sodium pyruvate, 0.292 g glutamine, 10 mL penicillin-streptomycin solution, 3.9 mg β-mercaptoethanol, 1 × 10⁵ U IL-2 (Biolegend), and RPMI-1640 to a final volume of 1 L. For acute stimulation, αCD3 antibody (Biolegend) was diluted to 2 μg/mL and added to a 24-well plate for overnight incubation at 4°C. The antibody solution was discarded the next day, and the wells were washed twice with pre-warmed PBS. A T cell suspension (1 × 10⁶ cells/mL) was added to the coated wells, and soluble αCD28 antibody (Biolegend) was added to a final concentration of 0.5 μg/mL. The plate was incubated at 37°C in a 5% CO₂ incubator for 48 hours. After 48 hours, cells were collected by gentle pipetting, transferred to a 1.5 mL microcentrifuge tube, centrifuged at 200 × g for 5 minutes, and the supernatant was discarded. The cells were resuspended in fresh medium containing 10 ng/mL IL-2 and continued to culture until the experiment endpoint. If the cell density exceeded 2 × 10⁶ cells/mL, fresh medium was added for dilution. For chronic stimulation, αCD3 antibody was diluted to 5 μg/mL and added to a 24-well plate for overnight incubation at 4°C. After washing twice with pre-warmed PBS, a T cell suspension was added, and soluble αCD28 antibody was added to a final concentration of 2 μg/mL. IL-2 was maintained at 10 ng/mL in the medium, and the plate was incubated at 37°C in a 5% CO₂ incubator for continuous culture. The coating plate and medium were replaced every 48 hours to ensure continuous antigen stimulation. For cell passage, cells were collected every 48 hours by gentle pipetting, transferred to a 1.5 mL microcentrifuge tube, centrifuged at 200 × g for 5 minutes, and washed with 1 mL of pre-warmed PBS. The cells were resuspended in fresh complete medium and cultured at 37°C in a 5% CO₂ incubator. The cell morphology and density were regularly monitored using an inverted microscope. For drug treatment, 3-IAA and GSK2606414 (GSK) were added to the culture medium at concentrations of 1 μg/mL and 1 μM, respectively, and the treatment was continued for 8 days following T cell activation.

### T cell in vitro proliferation assay

A 1 μM stock solution of carboxyfluorescein succinimidyl ester (CFSE, Thermo Fisher) was prepared in PBS and stored in the dark. T cells were isolated from mouse spleens and activated via acute stimulation as described previously, with or without 3-IAA treatment for the experimental groups. The isolated and washed T cells were resuspended in pre-warmed PBS (37°C) containing 0.1% bovine serum albumin (BSA) to a density of 1 × 10⁷ cells/mL to ensure uniform staining. CFSE working solution (1 μM) was prepared from the stock in PBS with 0.1% BSA, and 1 μL of 1 mM CFSE stock was added per 1 mL of the cell suspension. The cells were incubated at 37°C in the dark for 10 minutes, with gentle mixing every 3-4 minutes to prevent cell sedimentation and ensure proper binding of CFSE to the cell membrane. After incubation, staining was immediately terminated by adding five volumes of pre-chilled complete medium, followed by mixing and incubation on ice for 5 minutes to reduce metabolic activity and minimize non-specific staining. The cells were centrifuged at 300 × g for 5 minutes, and the supernatant was discarded. Cells were resuspended in pre-chilled PBS containing 0.1% BSA, centrifuged again at 300 × g for 5 minutes, and washed three more times to remove unbound CFSE. After the final wash, cells were resuspended in complete medium and counted using a hemocytometer to adjust the cell density to 1 × 10⁶ cells/mL. If residual CFSE fluorescence remained, additional washes were performed until the supernatant showed no visible fluorescence. The stained T cells were cultured at 37°C in a 5% CO₂ incubator for the specified duration, with cell density checked every 24 hours. If cell density exceeded 2 × 10⁶ cells/mL, fresh medium was added for dilution. After cultivation, cells were collected by gentle pipetting, transferred to a 1.5 mL microcentrifuge tube, and centrifuged at 300 × g for 5 minutes. The supernatant was discarded, and cells were resuspended in PBS containing 0.1% BSA for flow cytometry analysis. The CFSE fluorescence intensity decay was measured by flow cytometry, and the proliferation index was calculated using FlowJo software.

### T cell in vitro cytotoxicity assay

OT1 T cells were isolated from mouse spleens and activated as described above, with or without 3-IAA treatment for the experimental groups. MC38-OVA-luc target cells were resuspended in DMEM medium containing 10% FBS and adjusted to a density of 1 × 10⁵ cells/mL. A 100 μL aliquot was added to each well of a 96-well flat-bottom plate. The plate was incubated at 37°C in a 5% CO₂ incubator for 6 hours to allow cell adhesion. Activated OT1 T cells were added to the target cell wells at the predetermined effector-to-target ratio, mixed, and incubated at 37°C in a 5% CO₂ incubator for 24 hours, with gentle shaking every 12 hours to ensure even cell contact. After co-culture, the supernatant was collected by centrifugation, and 50 μL of supernatant from each well was transferred to a fresh 96-well plate. LDH release was assessed using a commercial LDH assay kit (Beyotime) according to the manufacturer’s instructions. Briefly, 50 μL of supernatant was added to 50 μL of substrate mixture, mixed gently, and incubated in the dark at room temperature for 30 minutes, followed by the addition of 50 μL of stop solution. Absorbance was measured using a microplate reader. The specific lysis rate was calculated using the formula: (experimental group absorbance - control group absorbance) / (maximum absorbance - control group absorbance) × 100%.

### Immunohistochemistry

Paraffin-embedded tissue sections were deparaffinized by sequential immersion in xylene (twice, 15 minutes each), followed by rehydration through graded ethanol solutions (100%, 95%, 85%, and 75%, 5 minutes each), and washed in distilled water until neutral. Antigen retrieval was performed by microwaving the hydrated slides in citrate buffer (pH 6.0) until boiling, then maintaining at a sub-boiling temperature for 10 minutes, followed by cooling at room temperature for 20 minutes. Endogenous peroxidase activity was blocked by incubating the slides in 3% hydrogen peroxide solution at room temperature for 25 minutes, then washed in PBS (three times, 5 minutes each). Non-specific binding was minimized by incubating the sections with 3% BSA in PBS for 30 minutes at room temperature. Primary antibodies (CD8, Servicebio, GB12068, 1:1500) were applied and incubated overnight at 4°C. After washing in PBS (three times, 5 minutes each), biotinylated secondary antibodies were applied and incubated at room temperature for 50 minutes. Following additional PBS washes, slides were incubated with HRP-conjugated streptavidin at room temperature for 30 minutes. Color development was achieved by applying DAB substrate solution, monitoring under a microscope until the desired intensity was reached, and then rinsing with tap water. Nuclear counterstaining was performed with hematoxylin for 3 minutes, followed by differentiation and bluing treatments. Slides were dehydrated through graded ethanol solutions, cleared in xylene, and mounted with neutral balsam. Staining results were independently evaluated by two pathologists.

### Public data analysis

The pan-cancer gene expression and patient annotation datasets from The Cancer Genome Atlas The Cancer Genome Atlas (TCGA) were obtained from the Genomic Data Commons (GDC) of the National Cancer Institute(Thorsson et al., 2018). For the MedDiet Score, highly expressed genes in the MedDiet group (fold change > 2, p < 0.05) were identified in mouse models using the limma package, as detailed in Table S3, and these genes were employed to construct the MedDiet Score. Gene sets associated with T cell functional activity were sourced from Miao *et al*.(Miao et al., 2020), and the Gene Set Variation Analysis (GSVA) algorithm(Hänzelmann et al., 2013) was applied to compute sample-specific gene set activity scores across various pathways. To assess immunotherapy efficacy, the Cancer Immunology Data Engine (CIDE)(Gong et al., 2025) was utilized to evaluate the association between the MedDiet Score and clinical outcomes following immunotherapy in patient cohorts with melanoma(Gide et al., 2019; Liu et al., 2019; Riaz et al., 2017), lung cancer(Patil et al., 2022; Roper et al., 2021), urothelial carcinoma(Hamidi et al., 2024), hepatocellular carcinoma(Finn et al., 2020), renal cell carcinoma(McDermott et al., 2018), pancreatic cancer(Padrón et al., 2022), head and neck cancer(Foy et al., 2022), and mixed cancers(Li et al., 2023). For the relationship between 3-IAA and tumor prognosis, clinical outcomes and serum 3-IAA concentrations in the Hamburg and Munich pancreatic cancer patient cohorts were derived from Tintelnot *et al*.(Tintelnot et al., 2023).

### Statistical analyses

Data were analyzed using GraphPad Prism 8.0 and R software version 4.0.0. All experiments were performed with at least three independent biological replicates. For comparisons between two groups, data were assessed for normality; parametric Student’s t-test was applied to normally distributed data, while non-parametric Mann-Whitney U test was used for non-normally distributed data. For multiple group comparisons, one-way analysis of variance (ANOVA) followed by Tukey’s post hoc test was employed. Correlation analyses were conducted using Spearman’s correlation test. Survival curves were constructed using the Kaplan-Meier method, and intergroup differences were evaluated with the log-rank test. Statistical significance was defined as p < 0.05.

## Abbreviations

3-IAA: indole-3-acetic acid
ATCC: American Type Culture Collection
ATF4: activating transcription factor 4
B. thetaiotaomicron: Bacteroides thetaiotaomicron
BHI: brain heart infusion
BSA: bovine serum albumin
CFSE: carboxyfluorescein succinimidyl ester
CHOP: C/EBP Homologous Protein
CIDE: Cancer Immunology Data Engine
E. coli: Escherichia coli
eIF2α: eukaryotic initiation factor 2α
GDC: Genomic Data Commons
GSEA: gene set enrichment analysis
GSVA: Gene Set Variation Analysis
HCAR2: hydroxycarboxylic acid receptor 2
HFD: high-fiber combined with fish oil diet
HMDB: Human Metabolome Database
HOPX: homeodomain-only protein homeobox
HPFS: Health Professionals Follow-Up Study
ICB: immune checkpoint blockade
IFNγ: interferon-γ
ISR: integrated stress response
LEfSe: linear discriminant analysis effect size
MedDiet: Mediterranean diet
MyD88: myeloid differentiation primary response protein 88
NHS: Nurses’ Health Study
PULs: polysaccharide utilization loci
SCD: stearoyl-CoA desaturase
TAM: tumor-associated macrophage
TCGA: The Cancer Genome Atlas
TLR4: Toll-like receptor 4
TME: tumor microenvironment

## Declarations

### Ethics approval

All animal experiments were approved by the Animal Ethics Committee of Renmin Hospital of Wuhan University, and were performed in accordance with the relevant guidelines.

### Consent for publication

Not applicable.

### Data Availability

All data generated or analyzed during this study are included in the manuscript and supporting files. The raw RNA-seq and metagenomic sequencing data have been deposited in the China National Center for Bioinformation (CNCB) under accession numbers CRA035878, CRA035890, and CRA035904.

### CRediT authorship contribution statement

Xin Yu: Writing – review & editing, Writing – original draft, Validation, Investigation. Wenge Li: Writing – review & editing, Writing – original draft, Validation, Investigation, Funding acquisition. Hongfang Feng: Writing – review & editing, Writing – original draft, Validation, Investigation. Zhiyu Li: Writing – review & editing, Validation, Investigation. Hongmei Zheng: Writing – review & editing, Validation, Investigation. Shengrong Sun: Writing – review & editing, Supervision, Project administration, Funding acquisition, Conceptualization. Juanjuan Li: Writing – review & editing, Supervision, Project administration, Funding acquisition, Conceptualization. Bei Li: Writing – review & editing, Supervision, Project administration, Funding acquisition, Conceptualization. Qi Wu: Writing – review & editing, Supervision, Project administration, Funding acquisition, Conceptualization.

## Acknowledgements

The authors gratefully acknowledge BioRender (https://www.biorender.com) for creating professional schematic illustrations. We also extend our appreciation to Bullet Edits for providing expert English language editing services.

---

Figure S1. Antitumor effect of MedDiet is dependent on adaptive immunity. Immune cell infiltration levels analyzed using the CIBERSORT algorithm based on RNA-seq data. (*P < 0.05; **P < 0.01; ***P < 0.001; ****P < 0.0001)

Figure S2. 3-IAA alleviates the ISR to inhibit CD8+ T cell exhaustion. (A) CCK-8 assay quantifying relative cell numbers of MC38 cells treated with 3-IAA. (B, C) CFSE staining assessing proliferation of CD8⁺ T cells treated with 3-IAA: (B) Proliferation index; (C) Representative plots.

Figure S3. Serum 3-IAA levels are associated with favorable clinical outcomes in pancreatic-cancer patients. (A–C) Hamburg cohort: (A) serum 3-IAA concentrations in responders versus non-responders; (B) PFS and (C) OS Kaplan–Meier curves for patients with high versus low serum 3-IAA. (D) Munich cohort: PFS Kaplan–Meier curve for patients with high versus low serum 3-IAA.

